# Egg-surface bacteria deter oviposition by the oriental fruit fly *Bactrocera dorsalis*

**DOI:** 10.1101/2020.04.22.054999

**Authors:** Huijing Li, Lu Ren, Mingxue Xie, Yang Gao, Muyang He, Babar Hassan, Yongyue Lu, Daifeng Cheng

## Abstract

Finding a suitable oviposition site is a challenging task for a gravid female fly, since the hatched maggots have limited mobility, making it difficult to find an alternative host. The oriental fruit fly, *Bactrocera dorsalis*, oviposits on many types of fruits. Maggots hatching in a fruit that is already occupied by conspecific worms will face food competition. Here, we showed that maggot-occupied fruits deter *B. dorsalis* oviposition and that this deterrence is based on the increased β-caryophyllene concentration in fruits. Using a combination of bacterial identification, volatile content quantification, and behavioural analyses, we demonstrated that the egg-surface bacteria of *B. dorsalis*, including *Providencia sp*. and *Klebsiella sp*., are responsible for this increase in the β-caryophyllene contents of host fruits. Our research shows a type of tritrophic interaction between microorganisms, insects, and insect hosts, which will provide considerable insight into the evolution of insect behavioural responses to volatile compounds.

## Introduction

In most insects, gravid females select an appropriate and specific site for laying eggs. For example, oviposition site selection in blowflies (Diptera: Calliphoridae) depends on the surface area or volume of carcasses ^1^, *Bombyx mori* moths (Lepidoptera: Bombycidae) prefer to oviposit on mulberry leaves or sites containing mulberry leaf volatiles ^2^, and *Anopheles gambiae* (Diptera: Culicidae) prefers oviposition sites in habitats with fewer competitors and predators ^3^. *Pieris napi* (Lepidoptera: Pieridae) prefer to oviposit on plants with eggs compared to egg-free host plants, independent of the initial egg density ^4^ Several hypotheses have been proposed to explain this phenomenon. Female insects seem to take many aspects into account to give their offspring the best chances of survival ^5,6^. The females select a unique oviposition site for maximizing the survival, growth, and reproductive potential of the offspring ^7^. Modification of offspring phenotype, proximity to a suitable habitat for the progeny, maintenance of natal philopatry, or indirect oviposition site choice via mating choice could affect insect behaviours concerning oviposition site selection ^8^.

The olfaction, gustation, and tactile sense of insects play an essential role in their search for oviposition sites ^9^ The shapes ^10^, colours ^11^, and volatiles ^12^ of oviposition sites may provide relevant cues. Volatiles can be produced by insect hosts, insect faeces, and some microorganisms. For example, the larval faeces of *Phthorimaea operculella* (Lepidoptera: Gelechiidae) are a crucial oviposition deterrent for conspecific females ^13^. Faeces produced by mango fruit flies deter co-species from laying eggs ^14^ Similarly, *Drosophila melanogaster* (Diptera: Drosophilidae) showed strong oviposition aversion in the presence of faeces from carnivorous mammals that was linked to the phenol produced by pathogenic bacteria ^15^. For *Anopheles* mosquitoes, volatile substances from larval habitats can induce oviposition at low concentrations but reduce oviposition at higher concentrations ^16^. Similarly, blood inoculated with bacteria isolated from screwworms induced oviposition in female screwworms compared with the control ^17^. Furthermore, studies have indicated that the oviposition behaviour of flies can be affected by the egg surface bacteria of flies. For example, proliferating bacterial symbionts on house fly eggs can affect oviposition behaviour of adult house flies, *Musca domestica* ^18^. Such effect was also observed in the black soldier fly, *Hermetia illucens* (L.)^19^. However, these studies didn’t involve living host tissues that is modified by the bacteria.

*Bactrocera dorsalis* (Hendel) (Diptera: Tephritidae) is a notorious pest species known for causing immense economic losses due to its infestation of many types of commercial fruits and vegetables. The damage from this pest is caused by the larvae that bore inside the fruits. Host volatiles are vital substances used by *B. dorsalis* to locate feeding and oviposition sites. For example, the 1-octen-3-ol, (E)-Ethyl-2-methyl-2-butenoate and benzothiazole identified in mango fruit were proven to be oviposition stimulants for this pest species ^20^ Furthermore, *B. dorsalis* responds differently to different stages or parts of the host fruits. *B. dorsalis* prefers fully ripe mango over unripe mango ^21^ and lays eggs at the top of mango fruits rather than at the bottom ^22^. A few studies have reported host marking behaviour in *B. dorsalis*. However, the host marking pheromones of *B. dorsalis* have not yet been identified. Zhao et al. (2014) reported that egg-surface chemicals can be used as host marking pheromones by *B. dorsalis* ^23^; however, Liu et al. (2018) argued that the eggs did not produce host marking pheromones but rather the larvae produced these pheromones after hatching ^24^. Here, we showed that an increased concentration of β-caryophyllene in the host fruits was the host marking pheromone of *B. dorsalis* and that this increase was induced by the bacteria adhered to the laid eggs. We have identified the chemical compound responsible for *B. dorsalis* host marking behaviour and identified the origin and biological significance of that compound.

## Results

### Oviposition-deterring effect of egg-infested fruits

To test whether gravid females of *B. dorsalis* avoid oviposition in fruits harbouring eggs, we tested their oviposition preference in a 2-choice assay. The flies were allowed to oviposit in either un-infested or egg-infested fruit purees. The results of the oviposition assay indicated that the time after eggs were laid in fruits was the critical factor inducing oviposition avoidance in *B. dorsalis*. Un-infested guava, orange, and mango fruits were more attractive to the flies for oviposition compared to fruits infested with eggs. However, oviposition deterrence depended on the amount of time that had passed after the eggs were laid in the fruits, and significantly fewer eggs were laid in the egg-infested fruits 48 h after the initial egg deposition compared to the control fruits (Fig. 1). These results indicate that the chemicals deter oviposition should not be presented by or on eggs but produced after 48h.

**Fig. 1.**
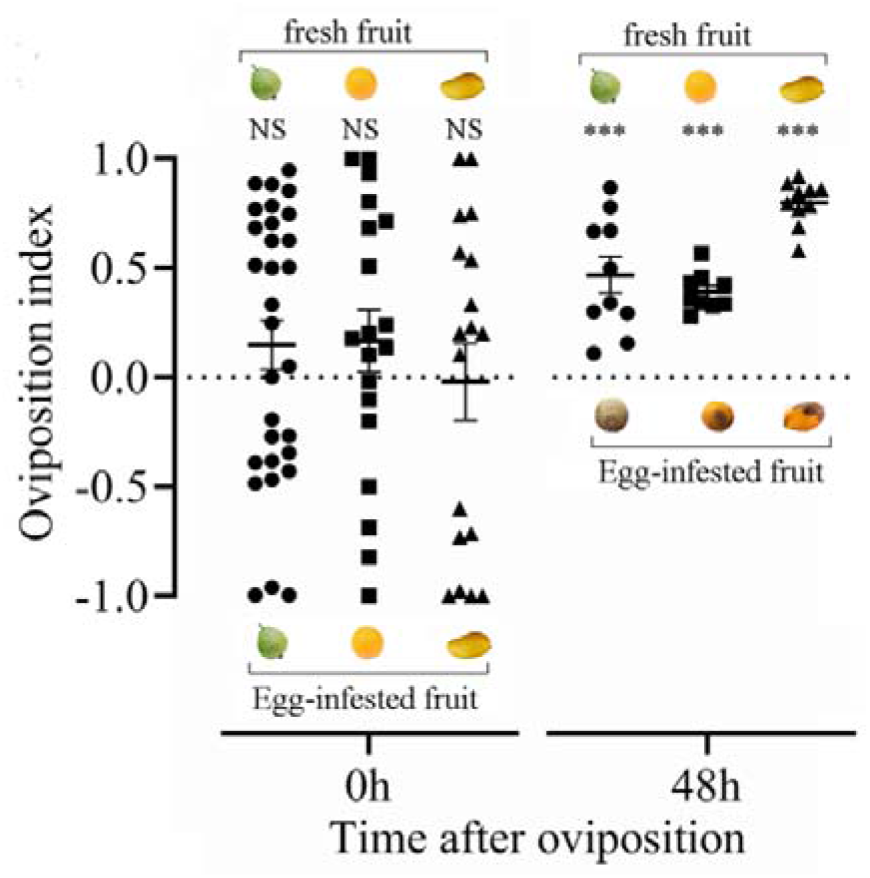
Oviposition index of *B. dorsalis* given a binary choice between fresh and egg-infested fruits. The circles, squares and triangles in the figure indicate the replicates. Error bars represent the SEM. The deviation of the response index from zero was analysed for significance with a paired sample Student’s t-test (NS: no significant difference, \*\*\**P* < 0.001).

### β-caryophyllene governs oviposition by *B. dorsalis*

To identify the active compound responsible for oviposition avoidance, we collected the volatiles produced by the egg-infested fruits using a solid-phase microextraction (SPME) fibre. Gas chromatography/mass spectrometry (GC/MS) analysis revealed that only α-copaene and β-caryophyllene were present in all fruits (different type of fruits or fruits with different treatments) in the experiments (Fig. 2A) which may indicate that α-copaene or β-caryophyllene plays a role in attracting or deterring the flies to lay eggs. However, comparison of the concentrations of α-copaene in the egg-infested and un-infested fruits (control) showed no significant differences among the two types of fruits (infested vs. un-infested) (Fig. 2B), while significantly higher levels of β-caryophyllene were detected in egg-infested fruits after 48 h compared to the control fruits (Fig. 2C). The average concentrations of β-caryophyllene were 150.8 ± 7.7, 105 ± 8.4 and 172.1 ± 11.5 μg/g in non-infested guava, orange, and mango, respectively, while the average concentrations of β-caryophyllene were 318.8 ± 52, 231.5 ± 31.7 and 331.3 ± 51.3 μg/g in egg-infested guava, orange, and mango, respectively. These results indicate that increased concentration of β-caryophyllene may play a role in deterring oviposition in egg-infested fruit.

**Fig. 2.**
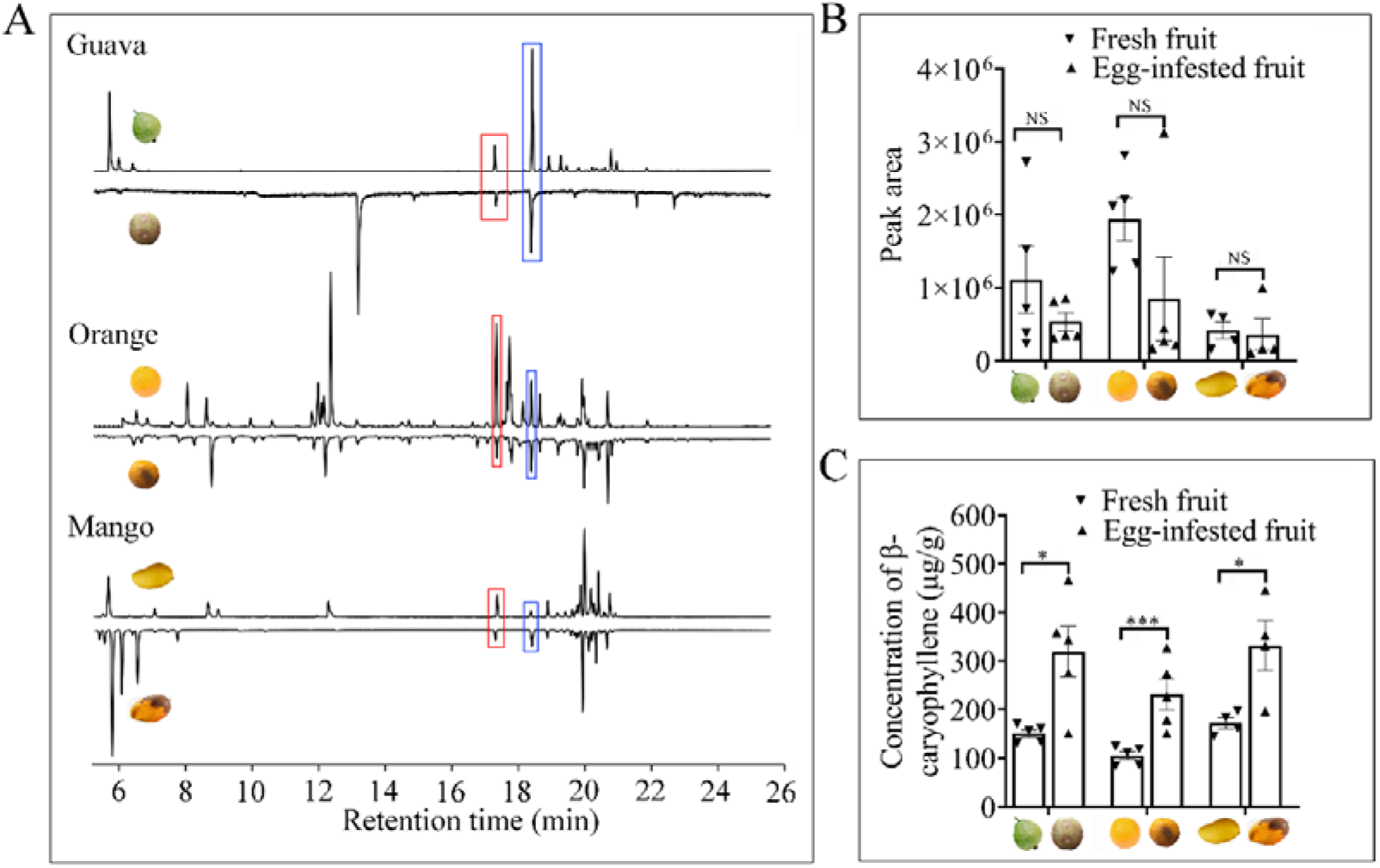
Gas chromatogram traces from odour collections of fruit samples. (A) Flame ionization (FID) traces from headspace collections of fresh fruits and fruits infested with eggs for 48 h. The red box indicates α-copaene, and the blue box indicates β-caryophyllene. (B) Peak area of α-copaene detected in fruits. (C) The concentration of β-caryophyllene in fruits. Error bars represent the SEM. The changes in α-copaene and β-caryophyllene contents were analysed for significance with independent sample Student’s t-test. (NS: no significant difference, \**P* < 0.05, \*\*\**P* < 0.001).

To investigate the impact of β-caryophyllene on *B. dorsalis* oviposition, we performed attraction assays to test the effect of different β-caryophyllene concentrations on the flies. In the four-arm olfactometer assay, the female flies were attracted at 150 μg/ml β-caryophyllene, which was almost the same as the concentration in fresh fruits (un-infested) (Fig. 3A). Moreover, The higher the concentrations of β-caryophyllene have the less attraction to the flies(Fig. 3A). GC-EAD measurements on female flies showed that all the females tested exhibited strong EAG responses to β-caryophyllene (Fig. 3B) which indicated that the olfactory sensilla in the antenna of *B. dorsalis* can be stimulated by β-caryophyllene. To address the behavioural sensitivity of *B. dorsalis* towards β-caryophyllene, we further tested oviposition preference in a 2-choice assay. Oviposition preference between fresh fruits and fruits to which β-caryophyllene was added showed that the flies significantly avoided laying eggs in the β-caryophyllene-amended fruits (Fig. 4). These results indicate that the different concentration of β-caryophyllene is the factor affects oviposition avoidance in female flies.

**Fig. 3.**
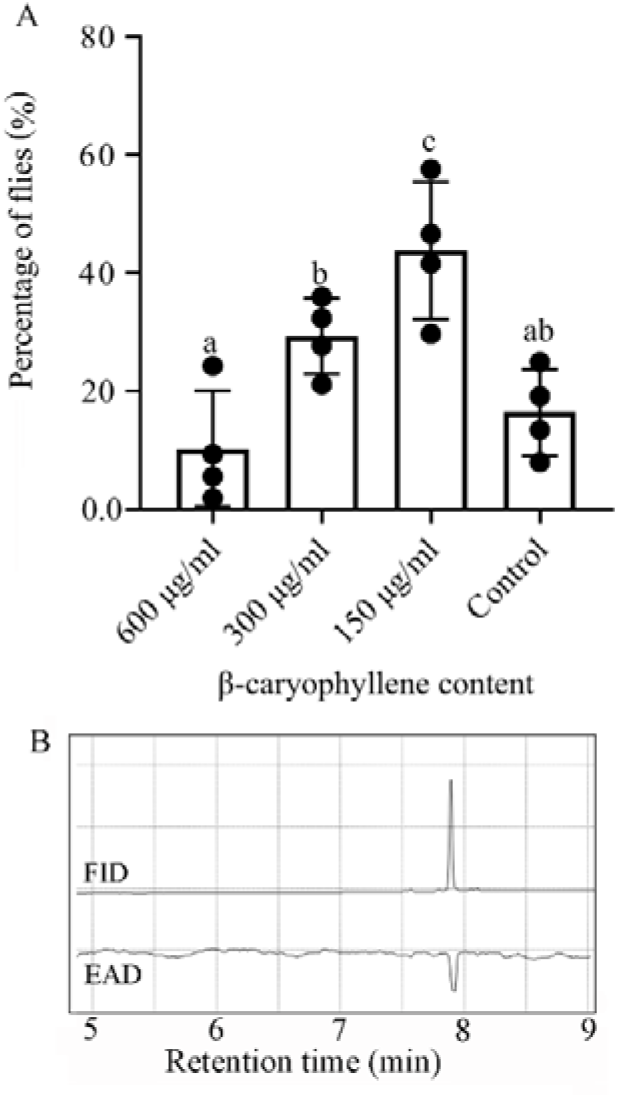
The response of female *B. dorsalis* to β-caryophyllene. (A) Choices of 100 females in a four-arm olfactometer. Different letters indicate significant differences among treatments at different concentrations by the Kendall nonparametric test at the 0.05 level. (B) Electroantennogram of female flies to 1 μl β-caryophyllene (150 μg/ml). FID: flame ionization detector; EAD: electrophysiological antennal detector (unit: mV).

**Fig. 4.**
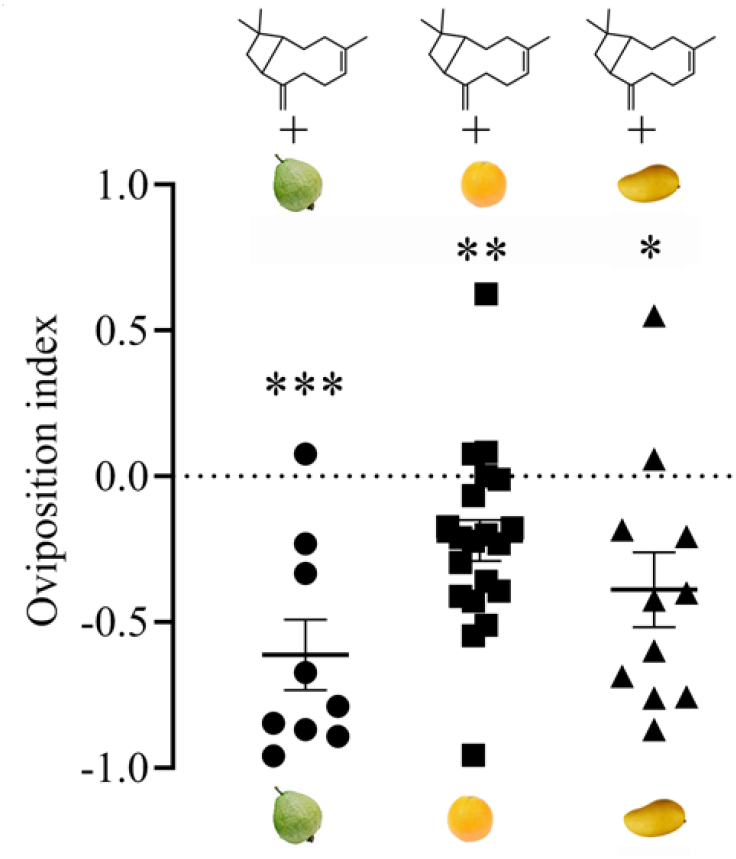
Oviposition index of *B. dorsalis* given a binary choice between fresh fruits and β-caryophyllene-amended fruits. The circles, squares and triangles in the figure indicate the replicates. Error bars represent the SEM. Deviation of the response index against zero was analysed for significance with paired sample Student’s t-test (\**P* < 0.05, \*\**P* < 0.01, \*\*\**P* < 0.001).

### Egg-surface bacteria increased the β-caryophyllene concentrations in fruits

Previous studies have indicated that some plants can release more β-caryophyllene after infection by some bacteria ^25–27^, and egg-surface bacteria can cause the fruit to rot ^28^. Thus, we hypothesized that egg-surface bacteria are involved in increasing β-caryophyllene in fruits. To test this hypothesis, we localized the bacteria present on the surfaces of fly eggs and tested the oviposition preference between fruits inoculated with egg eluent and control fruits. Using the Cy3-labelled general eubacterial probe, fluorescence in situ hybridization (FISH) visualization of the eggs and egg eluent revealed the presence of bacteria on the surfaces of eggs and in the egg eluent (Fig. 5A, B). β-caryophyllene concentrations in fruits inoculated with egg eluent were significantly increased compared to those in un-inoculated fruits (Fig. 5C), and the flies avoided laying eggs in fruits inoculated with egg eluent (after 48 h) (Fig. 5D). These results indicated that bacteria are presented on egg surface and might cause a change in oviposition preference.

**Fig. 5.**
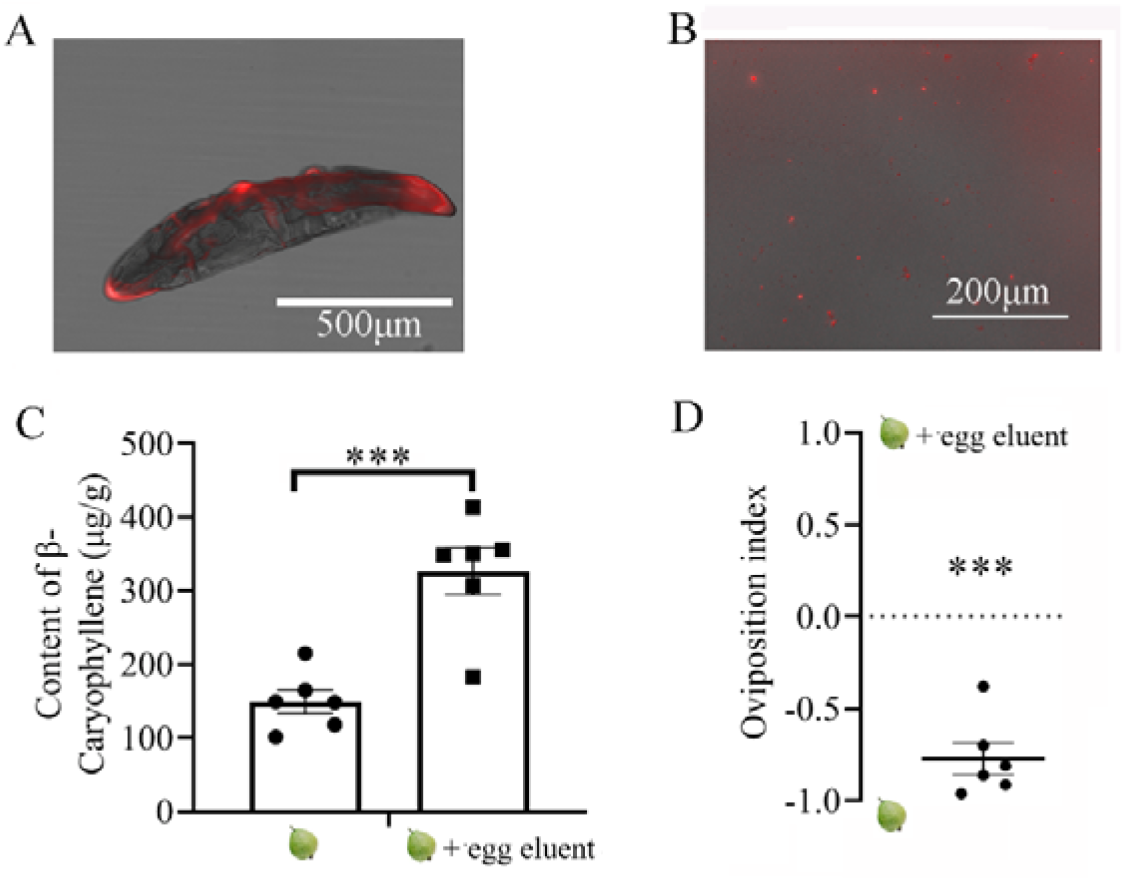
Effect of egg eluent on fruit and *B. dorsalis* oviposition behaviour. (A) Bacterial localization on the egg surface and in egg eluent (B) by FISH. Red signals indicate egg-surface bacteria. (C) Comparison of β-caryophyllene concentrations in fresh fruit and fruit inoculated with egg eluent after 48 h. Error bars represent the SEM. The change in the β-caryophyllene content was analysed for significance with paired sample Student’s t-test. (\*\*\**P* < 0.001). (D) Oviposition index of *B. dorsalis* given a binary choice between fresh fruits and fruits inoculated with egg eluent for 48 h. The circles in the figure indicate the replicates. Error bars represent the SEM. The deviation of the response index from zero was analysed for significance with a paired sample Student’s t-test (\*\*\**P* < 0.001).

### Isolation and identification of egg-surface bacteria

The identification and diversity of egg-surface bacteria was investigated using 16S rRNA gene sequencing. The results indicated that the dominant bacteria present on the surfaces of eggs belonged to *Providencia, Vagococcus, Salmonella, Lactococcus*, and *Klebsiella*, and *Providencia* accounted for more than 20% of the total bacterial abundance (Fig. 6A). Moreover, bacterial community profiles in egg-infested purees at different times indicated that there were significant differences in operational taxonomic units (OTUs) and Shannon diversity indexes (Fig. S1). A separation of the samples collected at different time points after inoculation with eggs was also shown by principal coordination analysis (PCoA) (Fig. S1). By comparing the differences in bacterial OTU abundance at different times, only the abundance of *Klebsiella* (in the five major egg surface genera in Fig. 6A) was found to be significantly increased after eight days (Fig. 6B). To verify whether egg-surface bacteria can affect the oviposition preference of *B. dorsalis*, the egg-surface bacteria were isolated. Three isolated bacterial strains were identified as *Providencia sp*. (accession number: MN894535), *Klebsiella sp*. (accession number: MN894536) and *Enterobacter sp*. (accession number: MN894537) (99% 16S rRNA sequence similarity) (Fig. 6C).

**Fig. 6.**
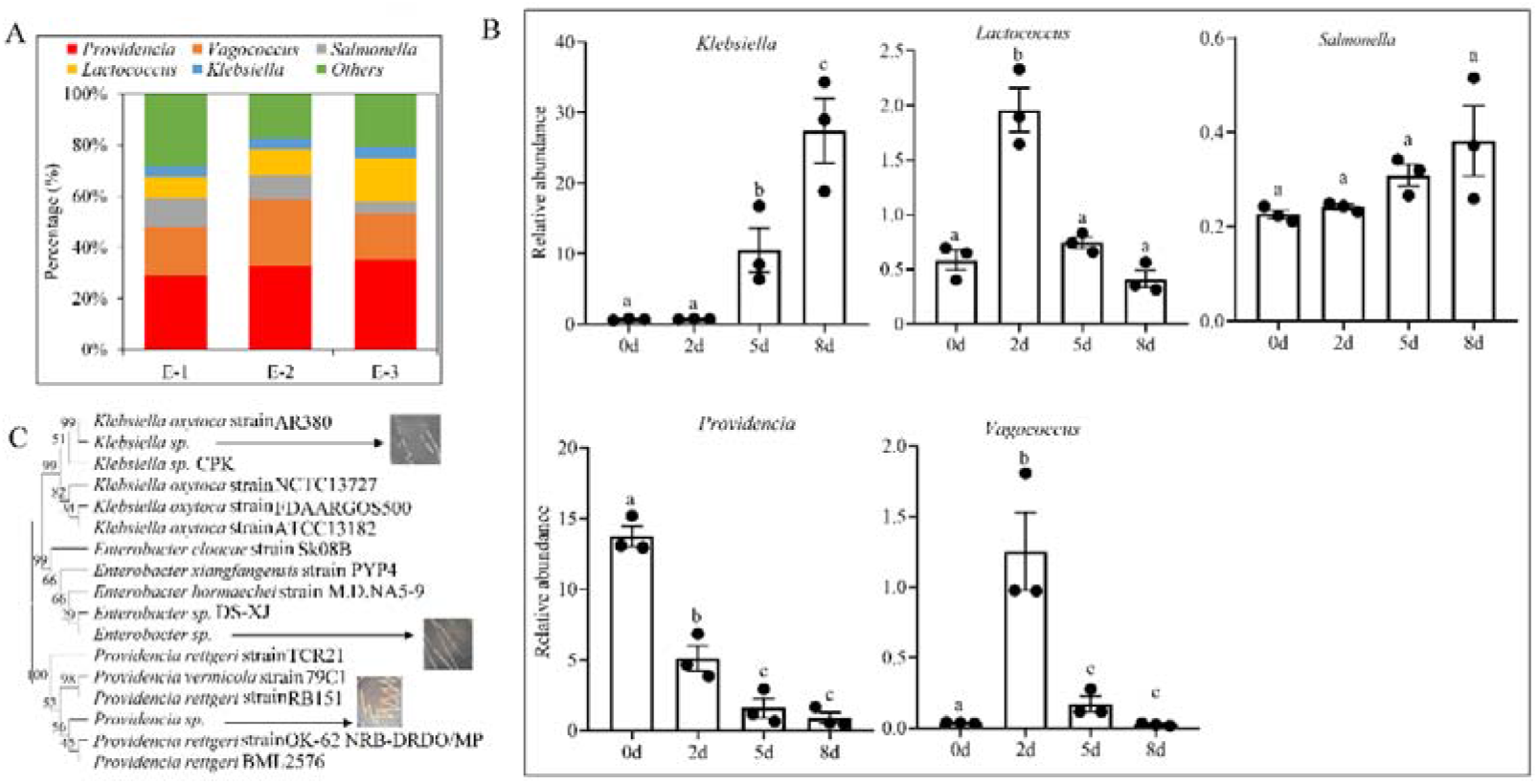
Egg-surface bacterial identification and isolation. (A) Egg-surface bacterial composition identified by 16S rRNA sequencing. (B) Relative abundance of egg surface bacteria in egg-infested fruits. Comparisons between groups were made using ANOVA followed by Tukey’s test. Significant differences (*P* < 0.05) are denoted by different letters. Error bars indicate the SEM. (C) Molecular phylogenetic analysis of egg-surface bacteria based on their 16S rRNA gene sequences. The tree was constructed with a maximum likelihood estimation method. Branch support is indicated by the bootstrap values (500 replicates).

### Specific egg-surface bacteria induced oviposition avoidance

To test the effect of the isolated bacteria on fly oviposition avoidance, guava fruits were inoculated with the isolated bacteria and then placed in oviposition cages. The oviposition selection assays showed that the flies avoided laying eggs in fruits inoculated with *Providencia sp*. and *Klebsiella sp*. (Fig. 7A), and the concentration of β-caryophyllene was significantly increased in these fruits (Fig. 7B). However, no significant differences in egg number and β-caryophyllene content were identified in fruits inoculated with *Enterobacter sp*. (Fig. 7). These results indicate that the egg-surface *Providencia sp*. and *Klebsiella sp*. can increase the concentration of β-caryophyllene in fruits, which can, in turn, lead to oviposition avoidance in female flies.

**Fig. 7.**
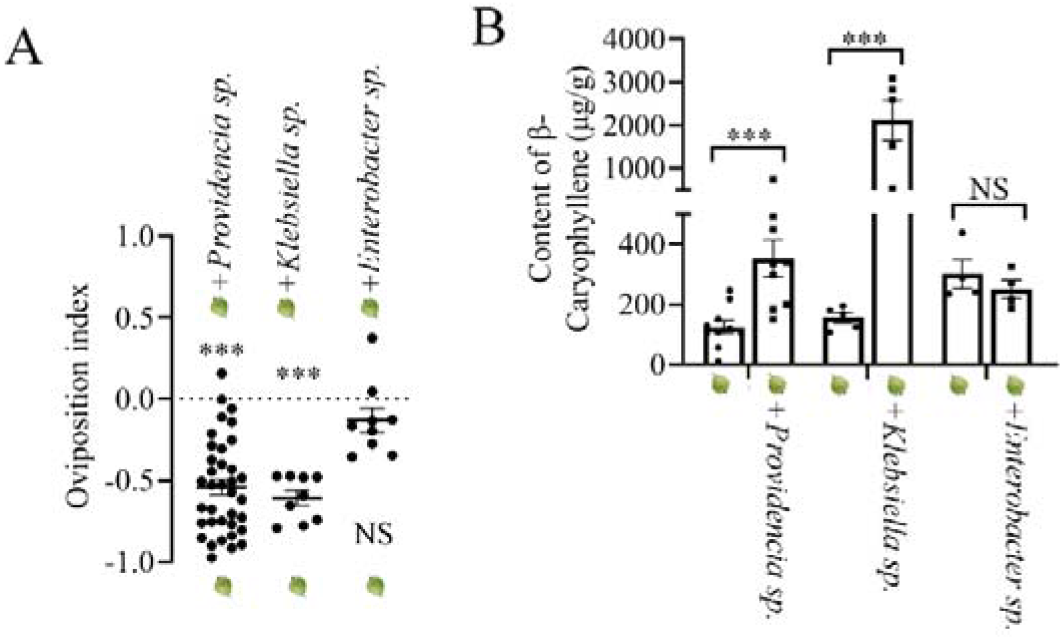
Effect of the isolated bacteria on the oviposition behaviour of *B. dorsalis*. (A) Oviposition index of *B. dorsalis* given a binary choice between fresh fruits and bacteria-inoculated fruits after 48 h. Error bars represent the SEM. The deviation of the response index from zero was analysed for significance with paired sample Student’s t-test (NS: no significant difference, \*\*\**P* < 0.001). (B) Comparison of β-caryophyllene concentrations between fresh fruit and fruit inoculated with bacteria after 48 h. Error bars represent the SEM. The change in the β-caryophyllene content was analysed for significance with a paired sample Student’s t-test. (NS: no significant difference, \*\*\**P* < 0.001). The circles AND squares in the figure indicate the replicates.

## Discussion

In this study, we investigated how the bacteria present on the surface of eggs can affect the oviposition preference of *B. dorsalis*. Our behaviour assay indicated that the egg-infested fruit induced oviposition avoidance in the flies, and this oviposition avoidance may be innate since females tested in our experiments were raised on an artificial diet and had no prior exposure to any fruits. Chemical analysis demonstrated that the concentration of β-caryophyllene was the critical factor eliciting oviposition avoidance. We confirmed that the relatively low level of β-caryophyllene (150.8 ~ 172.1 μg/g) in fruits could attract the flies to lay eggs, while the higher concentration of β-caryophyllene deterred oviposition. Our results also showed that egg-surface bacteria induced a higher level of β-caryophyllene in fruits after the eggs were laid. To the best of our knowledge, this is the first report showing that the chemical signals of host plants were induced by egg-surface bacteria and can, in turn, affect the oviposition behaviour of flies.

The presence of a conspecific brood (eggs or larvae) usually modifies the oviposition behaviour of phytophagous insects or entomophagous parasitoids ^29^ Females can help to reduce the level of competition faced by their offspring by avoiding laying eggs on or in hosts bearing a brood ^29^. Previous studies have indicated that avoidance of occupied hosts is typically mediated by cues and/or signals associated with the ovipositing females, offspring, or insect faeces ^29–33^. Although studies have indicated that chemical changes in host plants can be used as host marking pheromones by gravid females ^34,35^. such changes in plant compounds are associated with damage caused by oviposition or by tissue destruction by larvae or adults. Moreover, there are some claims about the induced systemic changes in plants that benefit plants or phytophagous insects, and the situation is further complicated by the finding that feeding by phytophagous insects causes systemic production of plant volatiles that attract natural enemies and thus may be favoured from the perspective of the host plant ^36–38^. In our study, we found that the host plant-produced β-caryophyllene was used as a host marking pheromone by *B. dorsalis*, and the bacteria present on the surfaces of eggs induced changes in the β-caryophyllene contents of plants. To our knowledge, this is the first time that a new mechanism of oviposition deterrence has been reported in insects. Although many studies have reported that plant volatiles can attract the natural enemies of *B. dorsalis* larvae ^39–42^. none of the volatiles were β-caryophyllene. Thus, the β-caryophyllene evolved by the host plants may favour the phytophagous insects in our situation. In addition, there are some studies have showed that egg surface bacteria can affect oviposition in insects^18,19^. However, bacteria referred in these studies didn’t involve living host tissue and the chemical cues that affect the oviposition behavor was not identified. With our results, the case that the effect of egg bacteria on oviposition choice may be widespread across insects, especially the cues that affect the oviposition behaviour in house fly depend upon a key bacterial strain *Klebsiella* ^18^.

We demonstrated that females responded differently to different concentrations of β-caryophyllene. Studies have revealed that the behavioural response of insects switches with changes in the concentrations of chemical cues. The responses of *Drosophila* to vinegar ^43^, ethyl acetate ^44^. and salt ^45^ are dose-dependent. We found that the relatively low concentration of β-caryophyllene can attract gravid females, while a higher level of β-caryophyllene deterred oviposition in the fruits, showing that the olfactory response of gravid females of *B. dorsalis* to β-caryophyllene is dose-dependent. The volatiles released by host plants provide essential information for phytophagous insects to identify and locate their hosts ^46–48^. β-caryophyllene is a typical terpenoid emitted by many kinds of plants, and β-caryophyllene plays an essential role in attracting or avoiding phytophagous insects. For example, the honeybee *Apis cerana* can use β-caryophyllene to locate its host flower, *Jacquemontia pentanthos* ^49^; β-caryophyllene also functions as a host-locating signal for the rice white-backed planthopper ^50^. However, other studies have indicated that β-caryophyllene can repel insects ^51–53^ or even be toxic to insects ^54^. Researchers have shown that *B. dorsalis* has oviposition preference for fruits at different developmental stages and identified the volatile profiles of the fruits ^21,55,56^ In these studies, we found that β-caryophyllene concentrations were higher in the fruits that the flies favoured for oviposition. However, the possible association between oviposition preferences and β-caryophyllene concentrations were not uncovered until this study.

Several studies have reported positive correlations between plant volatile terpenoid emissions and resistance to plant pathogens ^57^. For example, downy mildew *(Plasmopara viticola*)-resistant grapevine genotypes emitted significantly more monoterpenes and sesquiterpenes than susceptible genotypes ^58^. Rice genotypes resistant to the bacterial pathogen, *Xanthomonas oryzae* pv. *oryzae*, were shown to emit large quantities of either sesquiterpenes or monoterpenes ^59,60^. Both compounds were toxic to the bacterium at physiologically relevant concentrations *in vitro* ^59,60^ In citrus, higher emissions of C6 aldehydes (GLVs) and monoterpenes were also correlated with plant tolerance to huanglongbing disease ^61^. As a typical sesquiterpene, the content of β-caryophyllene in plants can also be induced by some bacteria ^25–27^. *Acinetobacter sp*. strain ALEB16 and *Gilmaniella sp*. strain AL12 can efficiently increase β-caryophyllene accumulation in *Atractylodes lancea* ^62^. Bacteria can activate signalling molecules, such as hydrogen peroxide (H2O2), salicylic acid, and jasmonic acid, which increase the expression of genes encoding key enzymes involved in β-caryophyllene biosynthesis pathways ^63^. For citrus, emission of β-caryophyllene increased significantly in both recently and long-term *Candidatus Liberibacter asiaticus*-infected citrus compared with controls ^64^. Similarly, apple trees infected by *Candidatus* Phytoplasma mali emitted more β-caryophyllene than healthy trees and were more attractive to a psyllid (*Cacopsylla picta*) vector ^65,66^. The increased β-caryophyllene in plants has demonstrated a role in plant defence against pathogens. For example, β-caryophyllene emitted from the stigmas of *Arabidopsis* flowers was shown to inhibit the growth of the pathogen *P. syringae* pv. *tomato* ^67^. From this point of view, the increased volatile terpenoids or β-caryophyllene in bacteria-infested plants may be a defence response of plants and may contribute to inhibiting bacterial infection. Thus, it is not surprising that the increased β-caryophyllene in *B. dorsalis* host fruits was induced by egg-surface bacteria and demonstrated a role in host fruit defence against egg-surface bacteria since surface bacteria can cause fruit rot ^28^.

Our results showed for the first time that the changed β-caryophyllene concentrations were sufficient to deter female *B. dorsalis* oviposition in a host fruit. We not only identified the responsible compound in the fruits but also revealed that egg-surface bacteria cause changes in β-caryophyllene concentrations in fruits. Hence, the current study increases the understanding of how animals cope with complicated chemical cues in the environment.

## Materials and methods

### Fly stocks

All experiments were carried out with a laboratory-reared *B. dorsalis* strain collected from a carambola (*Averrhoa carambola*) orchard located in Guangzhou, Guangdong Province, in April 2017. The flies were maintained in the laboratory under the following conditions: 16:8 h light:dark cycle; 70-80% relative humidity (RH); 25 ± 1 °C. A maize-based diet containing 150 g corn flour, 150 g banana, 0.6 g sodium benzoate, 30 g yeast, 30 g sucrose, 30 g paper towel, 1.2 ml hydrochloric acid and 300 ml water was used to feed the larvae, and the adult diet consisted of water, yeast hydrolysate, and sugar.

### Oviposition assays

The method adopted by Stensmyr et al. (2012) ^68^ was followed for the oviposition experiments. Briefly, a 2-choice apparatus was assembled in a cage made up of wood and wire gauze (length: width: height = 60 cm: 60 cm: 60 cm) with two Petri dishes (diameter: 3 cm) at the bottom of the cage (Fig. S2). All the devices were sterilized before each experiment. Then, the following tests were conducted to determine the oviposition preference of *B. dorsalis*.

The oviposition preference of the flies for non-infested or egg-infested fruits was tested using the following method (Fig. S3). Fresh fruits of guava (*Psidium guajava* Linn.), orange (*Citrus sinensis*), and mango (*Mangifera indica* L.) were sourced from the local market in Guangzhou, China. These fruits were sterilized the surface with ethanol (no bacteria can be detected inside with PCR detection) and then equally divided into two groups in a clean bench; one-half was put into a sterilized cage with 30 gravid female flies to inoculate eggs for 2h, the other half was also put into the cage but separated from the flies with a petri dish. Two hours later, the surface of the egg inoculated fruit was sterilized with ethanol (the presence of eggs in the fruits was confirmed by destructive sampling of a few fruits). Then, both groups of fruits were ground into puree with a sterilized grinder, and the puree (2 g) was added to the sterilized Petri dishes of the cages (one dish with puree containing eggs, one dish with puree without eggs). Thirty females were allowed to lay eggs for three hours in the puree. The number of eggs in each dish was counted, and the oviposition index was determined. Additionally, the oviposition preference of the flies to fresh fruit and egg-infested fruit (after 48 h) (larvae had hatched and the fruit had begun to rot) was also tested using the same procedure. Specifically, the fruit was removed after being laid eggs for 2 h and then the surface was sterilized with ethanol. Put the egg laid fruit and un-infested fruit into a sterilized cylindrical vessel and the vessel was put into the clean bench for 48h. For such experiments, the fruits will not be contaminated by the environmental bacteria since the fruits does not decay after 48h in control. For each assay, different flies were used to investigate oviposition preference. The numbers of biological replicates are shown in the figures.

The oviposition preference of flies for puree of fresh fruit (uninfested) and puree supplemented with β-caryophyllene was tested using fresh guava, orange, and mango fruits separately (Fig. S4). Two grams of puree of each fruit with or without β-caryophyllene (200 μg, the amount was referred to the concentrations in fresh and infested fruits) was added to the Petri dishes of the cage, and the oviposition index was calculated. The numbers of biological replicates are shown in the figures.

To test the oviposition preference of the flies for fresh fruit or fruit mixed with egg eluent, the fresh fruits of guava were sterilized with ethanol and equally divided into two halves in the clean bench (Fig. S5). Both halves were cut into smaller pieces (2×2 cm), and one-half of the pieces were mixed with 30 ml egg eluent. Pieces of the second half were mixed with 30 ml sterile water. Then the fruits were added into sterilized conical flasks (250 ml) that were placed on a shaker for 48 h at 25°C. For such experiments, the fruits will not be contaminated by the environmental bacteria since the fruits does not decay after 48h in control. For the preparation of egg eluents, eggs were collected using a sterilized Petri dish (Fig. S6) containing 2 ml orange juice (sterilized by high temperature). Specifically, orange juice was added in juice room as oviposition attractant, the egg will be laid into the egg collection room. Egg collection room and juice room were separated. Then eggs (1 g) were collected with a sterilized brush and washed with 30 ml sterile water. The same method was used to prepare egg eluent in the following assays. After 48 h, the fruits were ground into a puree, and 2 g of the puree was added to the Petri dishes inside the cages (one dish with puree inoculated with egg eluent, one with puree without egg eluent). Then, the oviposition index was calculated. The numbers of biological replicates are shown in the figures.

To test the oviposition preference of the flies for fresh fruit or fruit with egg-surface bacteria, fresh fruits of guava were equally divided into two halves and cut into smaller pieces as described above. Pieces from one-half were treated with 30 ml of egg-surface bacteria (*Providencia sp., Klebsiella sp*. and *Enterobacter sp*., 1×10^8^ cfu/ml) that was suspended in sterile water. The bacteria was isolated and identified from the egg surface, detail information was described in the following methods (Fig. S7). Pieces from the second half were treated with sterile water (30 ml) as the control treatment. All the sterilized treatments were done as described above and the fruits will not be contaminated by the environmental bacteria since the fruits does not decay after 48h in control. Forty-eight hours later, the fruits were ground into puree, and 2 g of the puree was added to the Petri dishes inside the cages (one dish with puree inoculated with egg-surface bacteria, one with puree without egg-surface bacteria). Then, the oviposition index was calculated. The numbers of biological replicates are shown in the figures.

The oviposition index was calculated using the following formula in all tests:

*Ovipostion index = (0 — C)/(O + C)*, where O is the number of eggs in the treatment dish, and C is the number of eggs in the control dish.

### Volatile analysis

The volatile organic compounds present in guava, mango, and orange fruits were analysed by gas chromatography-mass spectrometry (GC-MS). Briefly, 2 g of puree was added into a 20 ml bottle, and then a 100-μm polydimethylsiloxane (PDMS) SPME fibre (Supelco) was used to extract the headspace volatiles for 30 min. GC-MS was performed with an Agilent 7890B Series GC system coupled to a quadruple-type-mass-selective detector (Agilent 5977B; transfer line 250°C, source 230°C, ionization potential 70 eV). The fibre was inserted manually into the injector port (240°C) for desorption, and the volatiles were chromatographed on an HP-5 MS column (30 m, 0.25 mm internal diameter, 0.25 μm film thickness). Helium was used as the carrier gas at constant pressure (110.9 kPa). After fibre insertion, the column temperature was maintained at 50 °C for 1 min, increased by 5 °C/min to 140 °C and then increased by 10 °C/min to 250 °C, which was maintained for 10 min. β-caryophyllene identification was based on the comparison of the mass spectra of the samples with those listed in the NIST mass spectral library. Additionally, the identification of β-caryophyllene was confirmed by comparing its retention time and mass spectra with those of authentic standards purchased from suppliers. To determine the contents of Beta-caryophyllene in fruits, as related to increasing bacterial numbers, a standard curve was generated with the authentic standards of β-caryophyllene. The standard curves were prepared in triplicate (n=3) using 0.00225, 0.0225, and 0. 225, 2.25, 22.5, and 225 μg/μl solutions of β-caryophyllene (diluted by ethanol). The numbers of biological replicates are shown in the figures.

### Behavioural assays

Behavioural responses of *B. dorsalis* were evaluated using a four-arm olfactometer ^69^ (Fig. S8). A vacuum pressure pump pushed air through activated charcoal and an Erlenmeyer flask filled with distilled water, at a maintained airflow of 400 ml/min. Three concentrations of β-caryophyllene (0, 150, 300, and 600 μg/ml) were assessed in the behavioural test. One millilitre of volatile samples was added into the flavour bottle. This bottle was connected to the insect collection vial that was connected to the insect walking room. For each test, 100 flies were added to the insect walking room. After 10 h, the olfactometer was disassembled, and the numbers of flies in each insect collection vial were counted. The numbers of biological replicates are shown in the figures.

### GC-EAD analysis

GC-EAD analysis was performed to determine whether β-caryophyllene could elicit antennal responses to the mated females. The gas chromatograph (Agilent 7890A, USA) was equipped with an HP-5 capillary column (30 m□×□0.32 mm□×□0.25 μm, Agilent). The initial oven temperature was 40°C, which was increased to 260°C at 10°C/min and held for 15 min. Nitrogen was used as the carrier gas at 25 ml/min. One microliter of β-caryophyllene (150 μg/ml, according to the behaviour assay) was injected into the GC column (Splitless model) at 250°C with a flame ionization detector (FID) at 260°C. For EAD preparations, an antenna of a mated female was mounted between two microelectrodes with electrode gels. The antenna tip was cut slightly to facilitate electrical contact. Only one recording was made per antenna, with ten successful recordings performed in total. The signals from the antennae were analysed with GC-EAD 2012 software (version 4.6, Syntech).

### Visualization of egg-surface bacteria

FISH was used to visualize the bacteria on *B. dorsalis* eggs and in the egg eluent. FISH detection was performed following the method of Cheng et al. (2017)^70^ using the Cy3-labelled general eubacteria probe EUB338 (5’-GCTGCCTCCCGTAGGAGT-3’) ^71^. The newly laid eggs were incubated in a hybridization buffer [20 mM Tris-HCl (pH 8.0), 0.9 M NaCl, 0.01% sodium dodecyl sulfate (SDS), and 30% formamide] containing 50 nM probe. Following overnight incubation, the samples were thoroughly bathed in 1ml phosphate-buffered saline (PBS) for 3 min and mounted in SlowFade antifade solution (Molecular Probes, Chuo-ku, Japan). Then, the samples were observed under an epifluorescence microscope (Axiophot, Carl Zeiss, Shinjuku-ku, Japan).

The egg eluent samples were prepared by washing 1 g of eggs with 3 ml of sterile water (to concentrate bacteria). One microliter of egg eluent was smeared on a glass slide, and the glass slide was dried with an alcohol lamp. Then, the FISH procedure was the same as that described for the eggs. Specifically, after hybridization, the samples were washed with 1ml flow of PBS for three times.

### Egg-surface bacterial identification

To analyse the egg surface bacteria diversity, egg eluent was prepared by washing 5 g of eggs with 15 ml sterile water (to make sure that enough egg surface bacteria can be collected) (three replicate samples were prepared). Then, the bacteria in the eluent were collected by centrifugation (4000 RPM for 10 min), and the bacterial DNA was extracted using the Bacterial Genomic DNA Extraction Kit (Tiangen, Beijing, China, http://www.tiangen.com/asset/imsupload/up0250002001571219042.pdf) according to the manufacturer’s protocols. To analyse bacteria diversity in fruits infected by flies for different time, bacterial DNA was extracted from eggs laid in guava fruits for 0, 2, 5 and 8 days, respectively (Fig. S8). Briefly, surface sterilized guava fruits were placed for two hours in a sterilized cages with 30 gravid females to inoculate eggs. After the fruits were laid eggs, the surface was sterilized by ethanol and put into sterilized cylindrical vessels and the vessels were put into the clean bench. Then, the pureed samples (larvae and eggs in puree were removed) were collected at 0, 2, 5 and 8 days, respectively. Since no bacteria can be detected by PCR in the fresh fruits, no negative (egg-free) control was set. Then the bacterial DNA in the puree was extracted using the Bacterial Genomic DNA Extraction Kit (Tiangen, Beijing, China, http://www.tiangen.com/asset/imsupload/up0250002001571219042.pdf). To analyse the bacterial diversity, the 16S rRNA V3-V4 region was amplified with PCR (95°C for 2 min, followed by 27 cycles at 98°C for 10 s, 62°C for 30 s, and 68°C for 30 s and a final extension at 68°C for 10 min) using the primers 341F: 5’-CCTACGGGNGGCWGCAG-3’ and 806R: 5’-GGACTACHVGGGTATCTAAT-3’. PCR was performed in triplicate in 50 μl mixtures containing 5 μl of 10 × KOD Buffer, 5 μl of 2.5 mM dNTPs, 1.5 μl of each primer (5 μM), 1 μl of KOD Polymerase, and 100 ng of template DNA. Amplicons were extracted from 2% agarose gels and purified using the AxyPrep DNA Gel Extraction Kit (Axygen Biosciences, Union City, CA, U.S.) according to the manufacturer’s instructions and quantified using QuantiFluor-ST (Promega, U.S.). Purified amplicons were pooled in equimolar concentrations and paired-end sequenced (2 × 250) on an Illumina Hiseq2500 platform according to standard protocols. Raw reads were removed if they contained more than 10% unknown nucleotides (N) or fewer than 80% of bases with quality values (Q-values) > 20. Paired-end clean reads were merged as raw tags using FLASH (v 1.2.11) with a minimum overlap of 10 bp and mismatch error rate of 2%. Noisy sequences ^72^ of raw tags were filtered with the QIIME (V1.9.1) pipeline under specific filtering conditions ^73^ to obtain high-quality clean tags. Clean tags were searched against all the reference databases provided in UCHIME (http://drive5.com/uchime/uchime_download.html) to perform reference-based chimaera checking using the UCHIME algorithm (http://www.drive5.com/usearch/manual/uchimealgo.html). All chimeric tags were removed, and the remaining tags were subjected to further analysis. Tags were clustered into OTUs of□≥□97% similarity using the UPARSE pipeline ^74^ The tag sequence with the highest abundance was selected as a representative sequence within each cluster. The representative sequences were classified into organisms with a naive Bayesian model using the RDP classifier (version 2.2) ^75^ based on the SILVA database (https://www.arb-silva.de/). (Sequencing data were submitted to NCBI, accession number: PRJNA384497 and PRJNA623231).

### Microorganism isolation and identification

For egg-surface bacterial isolation, 1 ml of egg eluent (1g eggs in 3 ml) was added to 90 ml of sterile water and shaken for 20 min. Then, 1 ml of this liquid was added to 9 ml of sterile water and diluted to concentrations of 10^-5^ and 10^-6^. A 200 μL volume of the diluted liquid was then coated onto a Luria-Bertani (LB) plate and cultured for 1 day. Pure bacterial colonies were selected by subculturing on LB media and stored in a 25% glycerol solution at −80°C. A Bacterial Genomic DNA Extraction Kit (Tiangen, Beijing, China) was used to extract the DNA of the bacteria according to the manufacturer’s instructions. Universal primers (F: 5’-AGAGTTTCATCCTGGCTCAG-3’ and R: 5’-TACGGTTAXXTTGTTACGACTT-3’) were used to amplify the 16S rRNA. The PCR products were confirmed by electrophoresis on a 0.8% agarose gel, and the target PCR product was sequenced. The 16S rRNA sequence was BLAST searched against NR database in NCBI (https://blast.ncbi.nlm.nih.gov/Blast.cgi). Based on the hits (percent of identity >99.9%, E-value = 0.0) from the BLAST search, a phylogenetic tree (sequences were aligned with MUSCLE in MEGA 6.0) of the identified bacteria was generated with MEGA 6.0 software. The maximum likelihood method (substitution model: Tamura-Nei model) was used to construct a phylogenetic tree based on the 16S rRNA sequences, and the phylogenetic tree was evaluated with 500 bootstrap analysis.

### Statistical analysis

The oviposition preferences, α-copaene content and β-caryophyllene content were analysed with paired sample Student’s t-test. The abundance of bacteria and fly percentages in the four-arm olfactometer assay were analysed with one-way or three-way analysis of variance (ANOVA) with Tukey’s post hoc tests. All statistical analyses were performed in SPSS 19.0.

### Data availability

Sequencing data were submitted to NCBI, accession number: PRJNA384497 and PRJNA623231; *Providencia sp*. (accession number: MN894535), *Klebsiella sp*. (accession number: MN894536) and *Enterobacter sp*. (accession number: MN894537).

## Acknowledgements

The work was supported by the National Natural Science Foundation of China (No.31601693).

## Author contributions

H. L., L. R., M. X., Y. G. and M. H did the experiments; B. H., Y. L. and D. C. wrote the manuscript.

