## supplementary figures for "Egg-surface bacteria deter oviposition by the oriental fruit fly *Bactrocera dorsalis*"

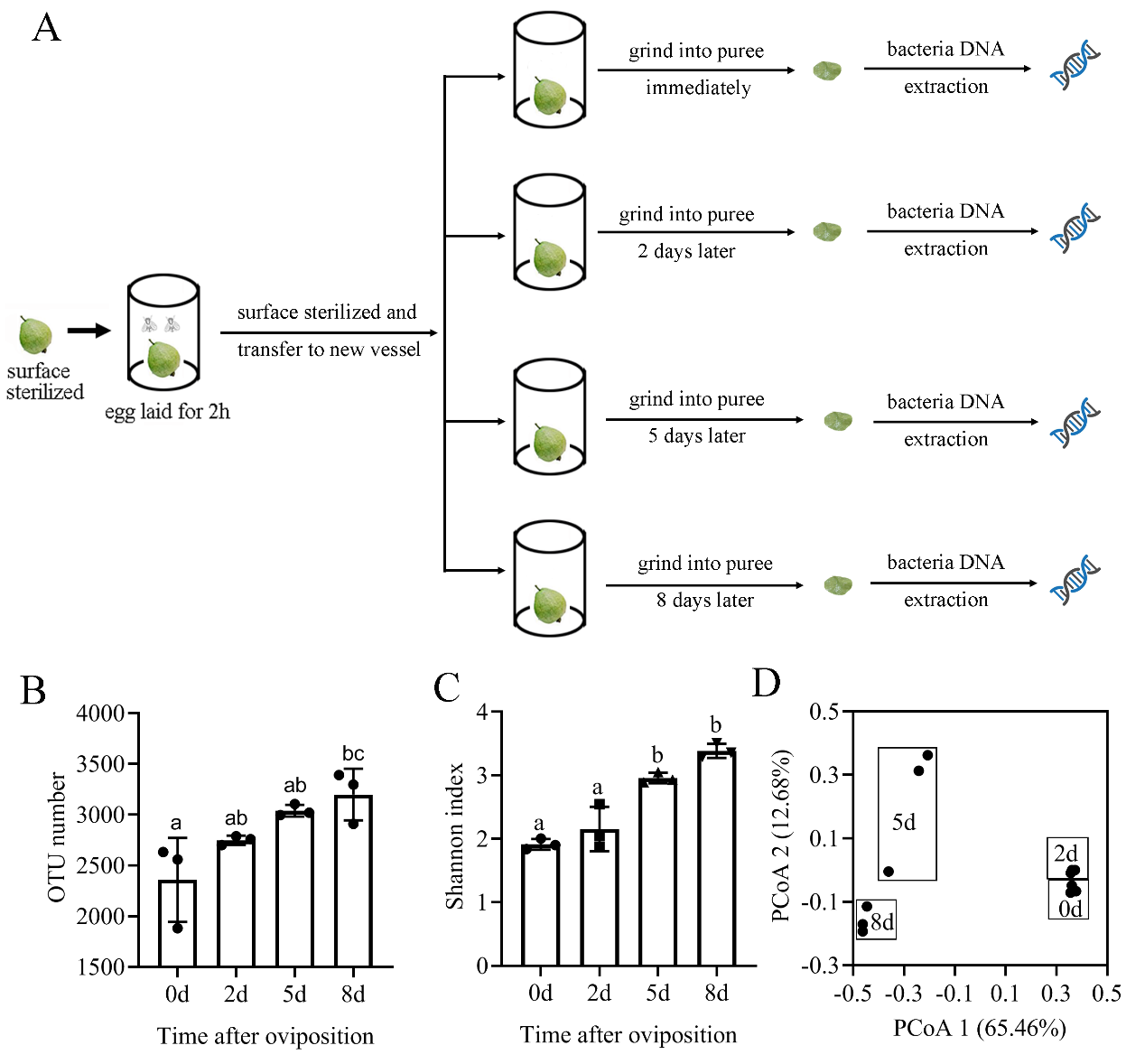

**Fig. S1.** Bacterial diversity in purees at different time intervals. (A) Experimental diagram for sampling. (B) OTUs and (C) Shannon diversity index values of the bacteria. (D) PCoA of the bacterial community in each sample. Different letters indicate significant differences in treatments by ANOVA at the 0.05 level.

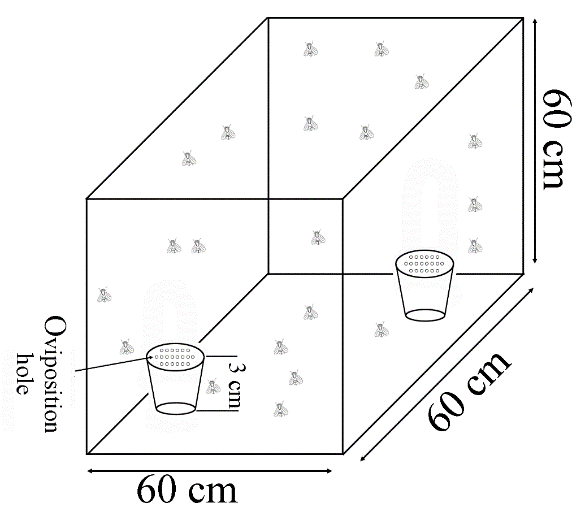

**Fig. S2.** Drawing of the 2-choice apparatus used to test the oviposition preferences of *B. dorsalis*.

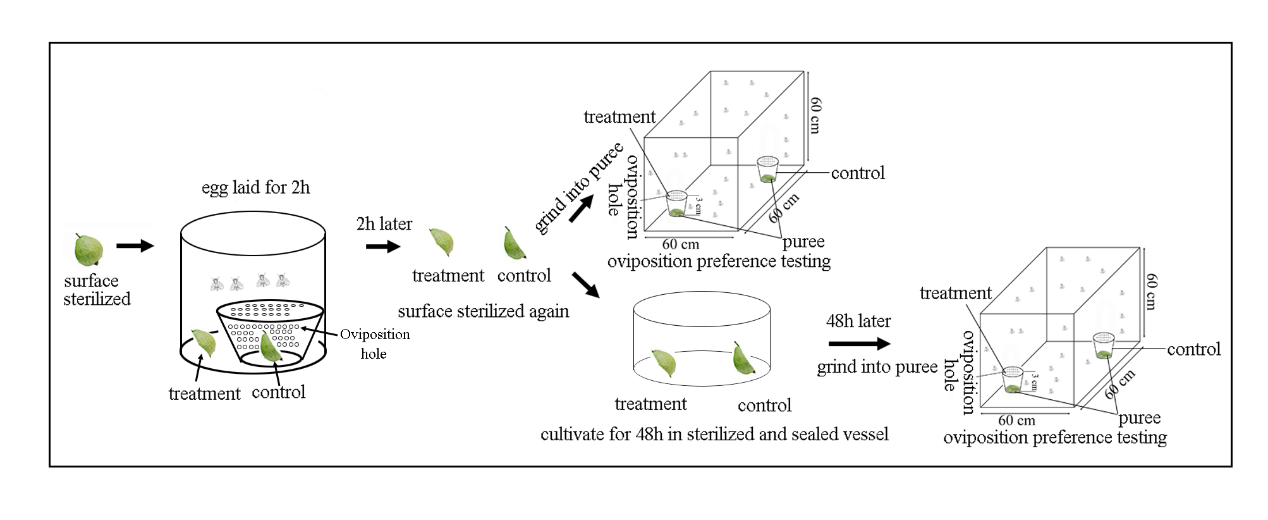

**Fig. S3.** Experimental diagram for testing the oviposition preference between fresh and egg-infested fruits.

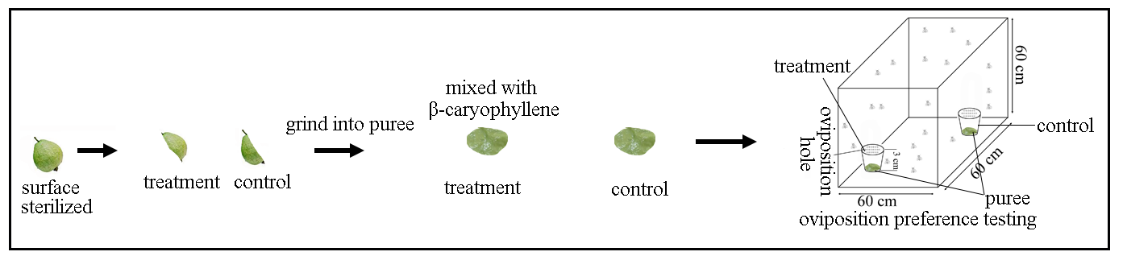

**Fig. S4.** Experimental diagram for testing the oviposition preference between fresh and β-caryophyllene-amended fruits.

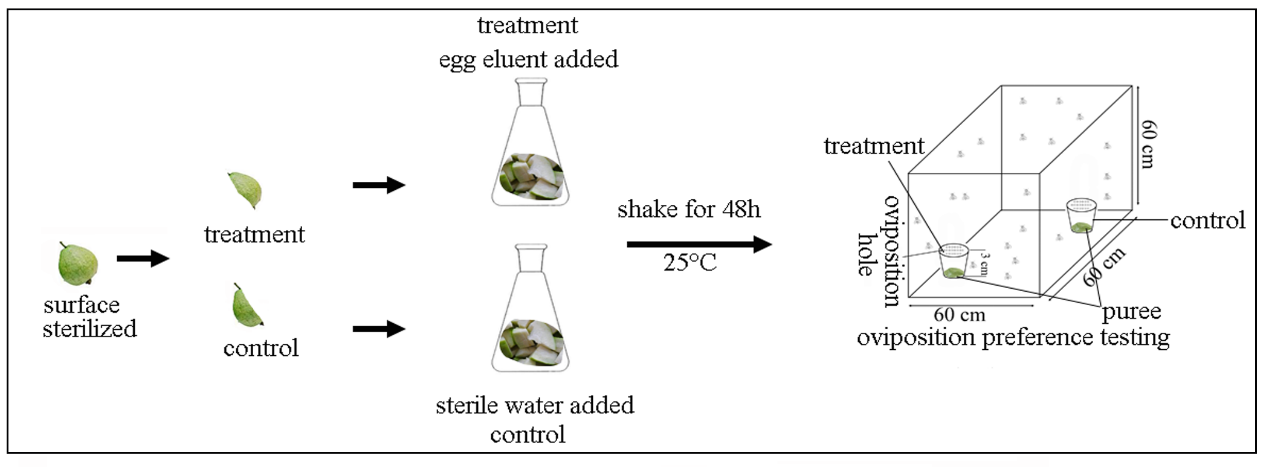
**Fig. S5.** Experimental diagram for testing the oviposition preference between fresh fruit and fruit inoculated with egg eluent after 48 h.

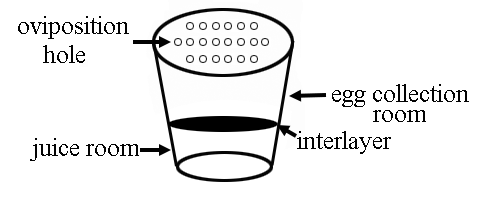

**Fig. S6.** Drawing of the petri dish used to collect eggs of *B. dorsalis*.

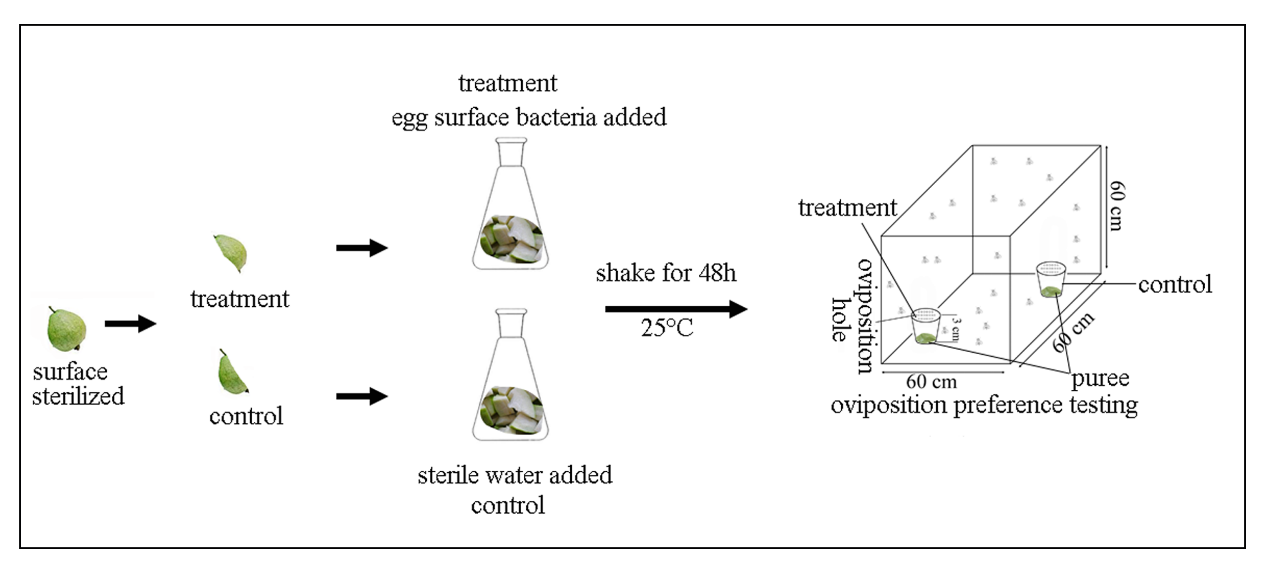
**Fig. S7.** Experimental diagram for testing the oviposition preference between fresh fruits and bacteria-inoculated fruits after 48 h.

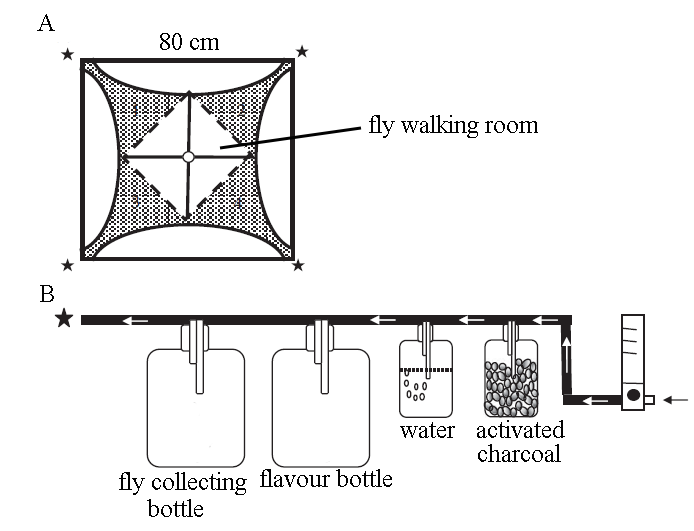

**Fig. S8.** Drawing of the four-arm olfactometer in which the attraction of flies to volatiles was tested. (A) Partition diagrams of the olfactometer: ★indicates the airflow inlet; ○indicates the central entrance for the fly; numbers (1, 2, 3, 4) indicate the final-choice area. (b) From right to left: the flow regulator, the activated charcoal bottle, the humidifier bottle, the flavour bottle and the fly collecting bottle.
